# A genome-wide CRISPR screen defines host determinants of early *Brucella* infection in human macrophage-like cells

**DOI:** 10.64898/2026.05.18.725962

**Authors:** Thomas Kim, Eleanor C. Scheeres, Aretha Fiebig, Andrew J. Olive, Sean Crosson

## Abstract

*Brucella* spp. are widespread intracellular animal pathogens that cause brucellosis, a significant zoonosis. Despite the global impact of brucellosis on animal and human health, the host genes that support *Brucella* infection remain incompletely defined. To address this knowledge gap, we developed a flow cytometry-based infection assay with fluorescent *Brucella* and performed a genome-wide CRISPR-Cas9 loss-of-function screen in human macrophage-like cells. Disruption of >150 host genes significantly reduced intracellular *B. abortus* burden at 3 h post-infection. In addition to recovering known host factors, the screen revealed previously unappreciated genes linked to endosomal trafficking, cytoskeletal remodeling, and lipid homeostasis. The screen was robust, as validation within these functional categories confirmed that the small GTPase RAB14, the Src-family kinase regulator CSK, and the phospholipid flippase subunit TMEM30A support early infection by *B. abortus* and *B. ovis* without impairing general phagocytosis. Gene set enrichment analysis further revealed positive regulators of mTORC1 signaling as key host factors; this result was validated through targeted disruption of LAMTOR2 and AKT1, and pharmacologic inhibition of AKT1. Thus, the AKT-Ragulator-mTORC1 signaling axis contributes to the establishment of a permissive intracellular niche during early *Brucella* infection. Finally, to assess whether these host requirements extend beyond *Brucella*, we examined infection by the unrelated intracellular pathogen *Mycobacterium abscessus*. CSK, AKT1, and LAMTOR2 were required for efficient *M. abscessus* infection, whereas RAB14 was dispensable. Together, these results define host genes that support early *Brucella* infection and distinguish shared versus pathogen-specific host dependencies exploited by intracellular bacteria.

## Introduction

*Brucella* spp. are intracellular pathogens that cause brucellosis, a globally distributed zoonosis that inflicts a significant burden on human and animal health (1, 2). Following host cell entry through actin-dependent mechanisms (3), *Brucella* localizes to a membrane-bound compartment known as the *Brucella*-containing vacuole (BCV), which undergoes a defined series of intracellular maturation steps. Newly formed BCVs initially traffic along the endocytic pathway, transiently acquiring early (Rab5, EEA1) and late (Rab7, LAMP1) endosomal markers while undergoing progressive acidification (4-7). Endosomal *Brucella*-containing vacuole (eBCV) acidification provides a cue that induces the VirB Type IV secretion system (T4SS), enabling delivery of bacterial effectors that remodel vesicular trafficking to support establishment of a replicative vacuole (rBCV) that is derived from the endoplasmic reticulum (8-12).

Multiple host surface molecules and membrane microdomains have been implicated in *Brucella* uptake, including sialic acid-containing receptors, components of the extracellular matrix (13), scavenger receptors (14, 15), complement receptors, cellular prion protein (16) and other lipid raft-associated molecules (17). Host complement proteins and antibodies can further modulate the route of entry and influence infection outcomes (17-21). These results indicate that *Brucella* infection mechanisms will vary depending on the host cell type, host exposure history, and the physiological state of the bacterium. Given the complexity and context-dependence of *Brucella* entry, a more integrated understanding of the host determinants that facilitate infection is needed.

Genetic approaches have begun to address this gap. For example, RNA interference (RNAi)–based screens in both invertebrate (22) and mammalian (23) host cells have highlighted roles for endoplasmic reticulum (ER) stress responses, autophagy, and vesicle trafficking in successful *Brucella* infections. Yet many questions remain regarding the host pathways that are required for *Brucella* to initiate a productive eBCV.

To build on this prior work, we conducted a genome-wide CRISPR–Cas9 loss-of-function screen to identify host genes required for *Brucella* infection of macrophage-like cells. Using a pooled knockout library in THP-1 cells and a flow cytometric readout of infection with highly fluorescent *Brucella* strains, we distinguished infected from uninfected cells by fluorescence-activated cell sorting (FACS) and quantified guide RNA representation between these populations by amplicon sequencing. Our results provide a genome scale framework for understanding early stages of *Brucella* infection of human macrophage-like cells. This study confirms several known host factors and identifies previously unrecognized host genes and pathways involved in endosomal trafficking, cytoskeletal remodeling, lipid metabolism, and mTOR signaling that support the early stages of mammalian cell infection by *Brucella* and other intracellular pathogens.

## Results

### Development of a flow cytometry-based approach to study early Brucella infection

To discriminate between infected and uninfected THP-1 macrophage-like cells by flow cytometry, we engineered highly fluorescent *Brucella* strains by integrating the mNeonGreen gene at the chromosomal *glmS* locus. We piloted this infection assay using *B. ovis*, a largely ovine-restricted pathogen that poses no zoonotic risk. Fluorescent *B. ovis* exhibited wild-type growth kinetics in axenic culture (Fig. 1A) and was readily visualized inside THP-1 cells by fluorescence microscopy. An MOI of 1,000 provided a clear and reproducible separation between infected and uninfected host cell populations at a 3 h timepoint (Fig. 1B, Fig. S1A). A shorter infection time (2 h) did not yield a sufficiently high enough number of infected cells to conduct a flow cytometry-based genetic screen (Fig. 1C, Fig. S1B). Importantly, there was no impact on THP-1 host viability following infection with *B. ovis* at an MOI of 1,000 for 3 h (Fig. 1D). We therefore proceeded to assess THP-1 infection by *B. abortus*::mNeonGreen using the same infection conditions. Flow cytometry showed that 50% of THP-1 cells were infected with *B. abortus* (Fig. 1E). We concluded that our infection protocol enabled sufficient *B. abortus* entry in a high enough percentage of cells to conduct a flow cytometry-based screen for host genes that impact the early stages of *B. abortus* infection.

**Figure 1.**
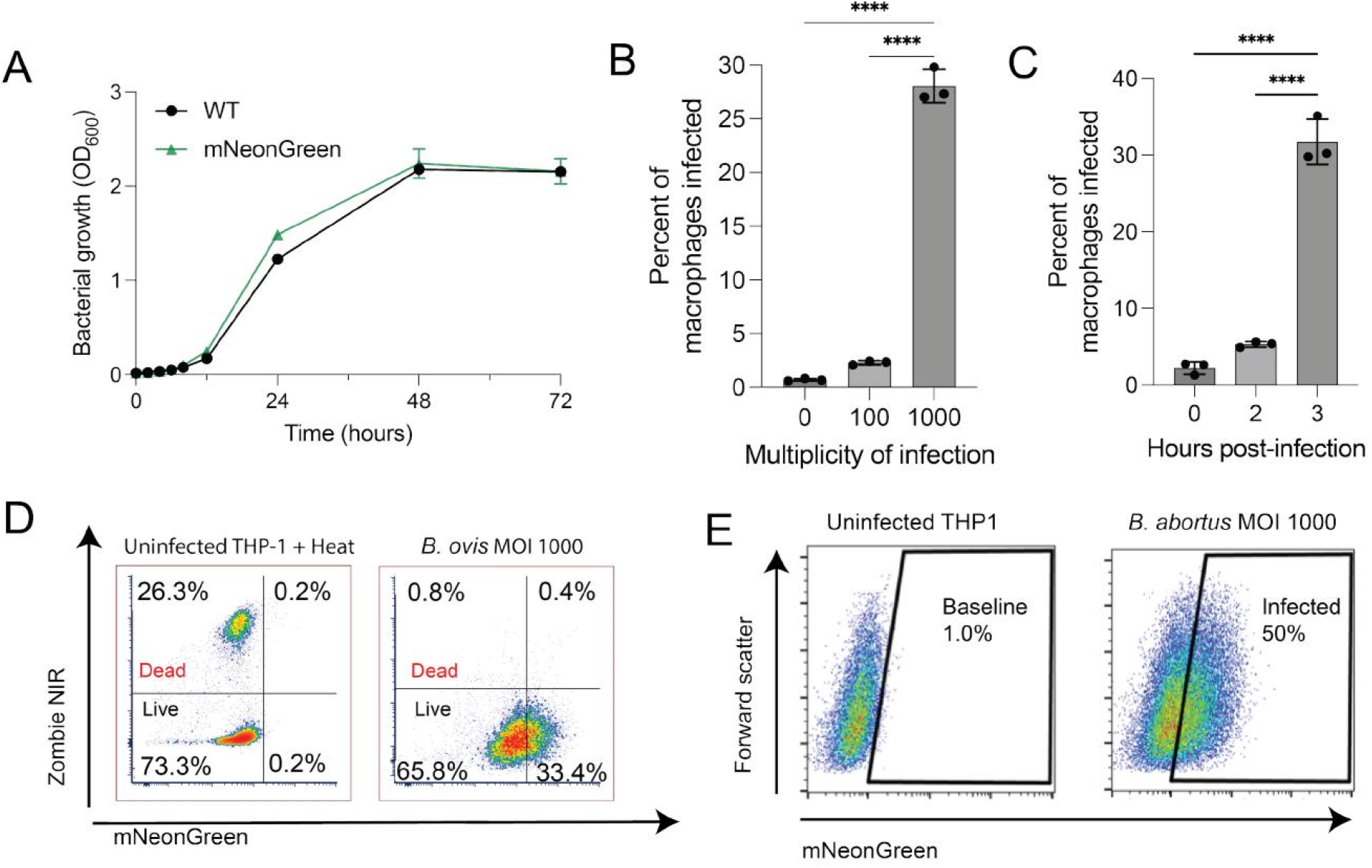
Flow cytometry enables detection of early *Brucella* infection in THP-1 macrophages. (A) Growth curves of wild-type (WT) and mNeonGreen-expressing *B. ovis* in rich medium over 72 hours post-inoculation. (B) Percentage of THP-1 macrophages infected with mNeonGreen *B. ovis* after 3 hours of infection at multiplicities of infection (MOIs) of 100 or 1,000, as assessed by flow cytometry. (C) Percentage of THP-1 macrophages infected with mNeonGreen *B. ovis* after 2 or 3 hours of infection at MOI 1,000. (D) Representative flow cytometry plots gated on live, single THP-1 macrophages. Cells were stained with Zombie NIR viability dye and infected with mNeonGreen *B. ovis* (MOI 1,000). The left panel shows a control population of heat-treated, uninfected macrophages. Percentages of live/dead (Zombie NIR) and infected (mNeonGreen-positive) cells are indicated. (E) Representative flow cytometry plots of THP-1 macrophages infected with mNeonGreen *B. abortus* (MOI 1,000), showing a distinct population of mNeonGreen-positive infected cells. Data in panels B and C represent means ± standard deviation (SD) from three biological replicates. Statistical significance was determined by one-way ANOVA followed by Dunnett’s multiple comparison test (***, *P* < 0.001; ****, *P* < 0.0001).

### CRISPR-Cas9 loss-of-function screen identifies host genes required for Brucella infection of macrophages

We generated a genome-wide loss-of-function library in immortalized, Cas9-expressing THP-1 monocytes using a pooled lentiviral set of single-guide RNAs (sgRNAs) targeting human genes (24) (Fig. 2A; see Materials and Methods for details on library design and quality metrics). To identify host genes that facilitate *B. abortus* entry and establishment of the endosomal *Brucella*-containing vacuole (eBCV), we infected a PMA-differentiated THP-1 CRISPR library with fluorescently labeled *B. abortus*. At 3 h post-infection, host cells were separated by FACS into the top 10% mNeonGreen-positive (infected) and bottom 10% mNeonGreen-negative (uninfected) populations (Fig. 2A). Genomic DNA was extracted from both groups, and sgRNAs were PCR-amplified and sequenced. Relative sgRNA abundance between the two sorted populations was compared to identify candidate host genes that impacted *Brucella* infection.

**Figure 2.**
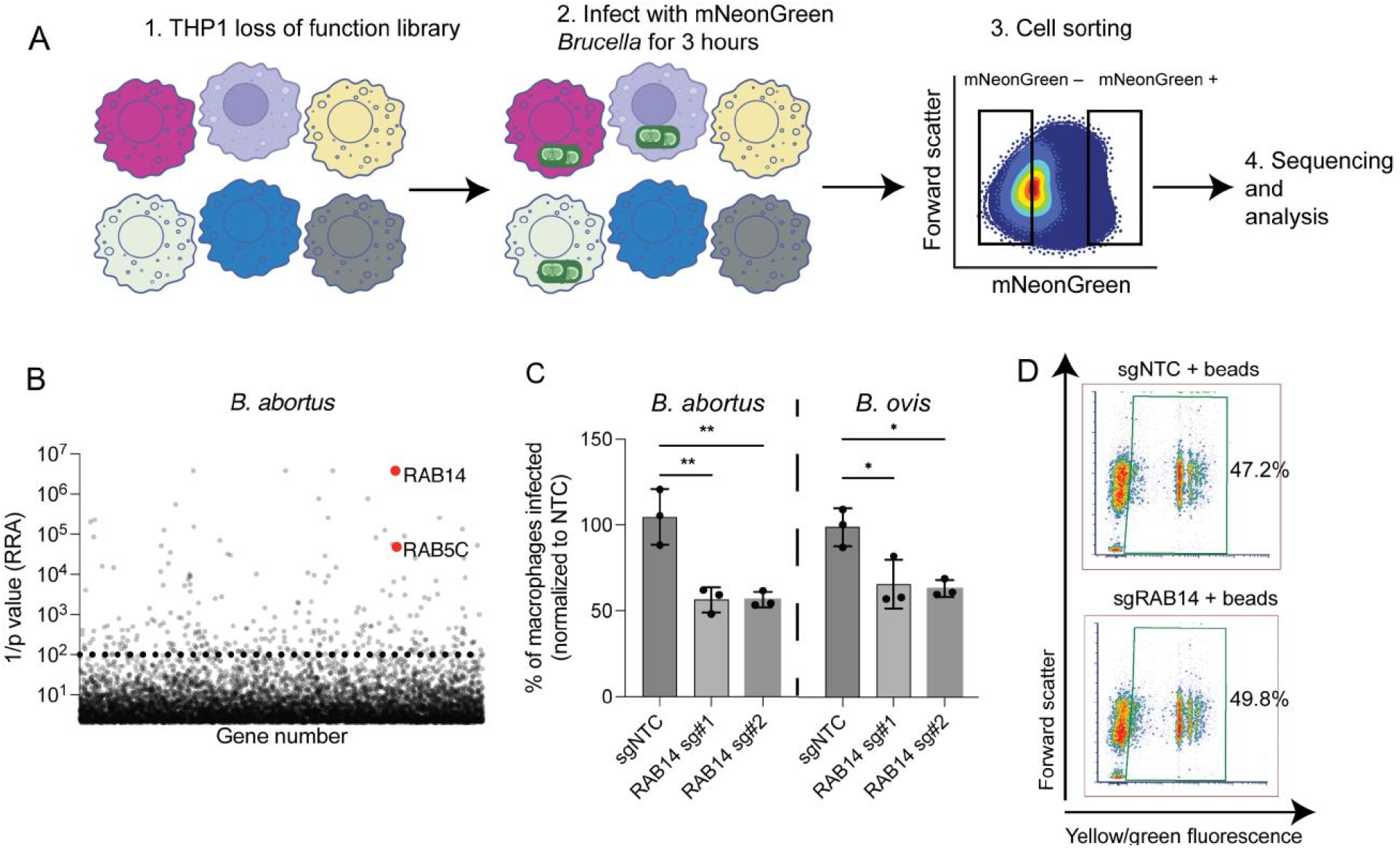
RAB14 is a host factor that facilitates *Brucella* infection of macrophages. (A) Schematic of the genome-wide loss-of-function screen. A CRISPR-Cas9 pooled sgRNA library was introduced into Cas9-expressing THP-1 macrophage-like cells, followed by infection with mNeonGreen-expressing *Brucella* for 3 hours. Infected (mNeonGreen^+^) and uninfected (mNeonGreen^−^) populations were separated by FACS, and the abundance of sgRNAs in each population was quantified by sequencing. (B) Plot of gene enrichment *P* values from the *B. abortus* CRISPR-Cas9 screen, calculated using the α-robust rank aggregation (α-RRA) algorithm in MAGeCK. *P*-value threshold of 10^-2^ (1/p of 10^2^) is drawn as a dotted line. The small GTPases *RAB14* and *RAB5C* are highlighted in red. (C) Validation of *RAB14* as a host factor. THP-1 macrophages expressing a *RAB14*-targeting sgRNA were infected with mNeonGreen *B. abortus* or *B. ovis* at MOI 1,000. The percentage of infected cells was quantified by flow cytometry and normalized to non-targeting control (NTC). Data represent means ± SD from three biological replicates. Statistical significance was determined by one-way ANOVA followed by Dunnett’s multiple comparison test (**, *P* < 0.01). Editing efficiency for both sgRNAs > 60% (D) Latex bead uptake assay in NTC and *RAB14*-deficient macrophages. Cells were incubated with yellow-green fluorescent beads (bead:cell ratio = 10) for 3 hours. Representative flow cytometry plots show comparable bead uptake between genotypes, indicating that phagocytic capacity is preserved in *RAB14*-deficient cells.

Host gene disruptions associated with increased intracellular *Brucella* burden were identified (Table S1), but gene set enrichment analysis (GSEA) of these putative restriction factors did not yield significant pathway-level enrichment. Furthermore, comparison of these putative restriction genes in our dataset to a previous siRNA screen of *B. abortus* infection in HeLa cells (23) revealed no overlap, with the exception of NME8. This limited concordance likely reflects differences in cell type (THP-1 macrophages versus HeLa epithelial cells), infection conditions, and readout timing, and suggests that restriction mechanisms may be more context- or cell type-specific than host-permissive pathways.

We therefore focused on host genes whose disruption significantly reduced intracellular *B. abortus* burden. Specifically, we identified over 150 host genes whose disruption was associated with reduced levels of *B. abortus* infection based on the following criteria: 1) detection of at least two sgRNAs in the dataset, 2) an experimental log selection ratio greater than 1, and 3) experimental *P* value less than 0.01 (Table S1). Several hits correspond to host factors previously implicated in *Brucella*-containing vacuole (BCV) biogenesis or maintenance, including calnexin (CANX) (10), CDC42 (3), and the early endosomal GTPase RAB5C (25, 26), underscoring the robustness of our screen and the biological relevance of our screen results. In addition, we identified CYFIP1, a core component of the WAVE regulatory complex that promotes actin polymerization at the plasma membrane (27); ARPC2, a subunit of the Arp2/3 complex that drives actin nucleation; and TBC1D10B, a GTPase-activating protein involved in endosomal recycling (28). In contrast to host restriction factors discussed above, these pathogen supportive genes were previously implicated in *B. abortus* infection in a prior siRNA screen in HeLa cells (23), lending support to our loss-of-function results.

### Endosomal trafficking and recycling genes supporting early Brucella infection: a role for RAB14

As described above, the endosome-associated small GTPase RAB5C and the GAP protein TBC1D10B were identified as host factors that promote early *B. abortus* infection (Table S1; Fig. 2B). Additional hits in genes involved in endosomal processes included ARF4, which regulates recycling endosome integrity (29) (Table S1). We further identified DNAJC13, a chaperone that alters the spatial organization of endosomal subdomains involved in recycling and degradation when disrupted (30, 31). A top hit in the *B. abortus* screen was the GTPase RAB14, which coordinates membrane trafficking at the interface of early and recycling endosomes and the trans-Golgi network (32-34) (Fig. 2B). To validate that RAB14 plays a role in infection, we generated RAB14-deficient THP-1 macrophages using two distinct sgRNAs and observed significantly reduced *B. abortus* intracellular burden compared to non-targeting control (NTC) cells (Fig. 2C; Fig. S2A). RAB14 disruption also diminished *B. ovis* infection (Fig. 2C), showing RAB14 to be a host factor for both the smooth zoonotic strain *B. abortus* and the naturally rough ovine-restricted pathogen *B. ovis*. To test whether the reduced *Brucella* infection observed in RAB14 mutant cells was due to a general defect in phagocytosis, we incubated the mutant THP-1 macrophages with fluorescent 1 µm latex beads. Bead uptake was comparable to non-targeting control cells (Fig. 2D), indicating that basic phagocytic capacity was intact. Thus, the infection defect in RAB14-deficient cells is not a result of impaired general phagocytosis.

### Expanding understanding of cytoskeleton regulatory genes that promote Brucella infection

Gene set enrichment analysis (GSEA) (35) of the ranked list of host mutants with reduced *B. abortus* infection revealed expected enrichment of functional categories involved in cytoskeletal and cell motility regulation (Table S1). Among these genes was C-terminal Src kinase (CSK) (Fig. 3A), a regulator that links G protein signaling to actin remodeling (36), and that has been implicated in dengue virus replication (37) and in the internalization of *Pseudomonas aeruginosa* by mammalian cells (38). To validate CSK as a host factor that supports *Brucella* infection, we generated a CSK mutant cell line. Infection assays using fluorescent *B. abortus* and *B. ovis* revealed significantly reduced mNeonGreen signal in the CSK mutant lines compared to control cells (Fig. 3B; Fig. S2B), supporting the conclusion that CSK facilitates infection by both *Brucella* species. Again, uptake of fluorescent 1 µm latex beads was comparable between mutant and control cells (Fig. 3C; Fig. S2D), indicating that basic phagocytic function was preserved in a CSK mutant.

**Figure 3.**
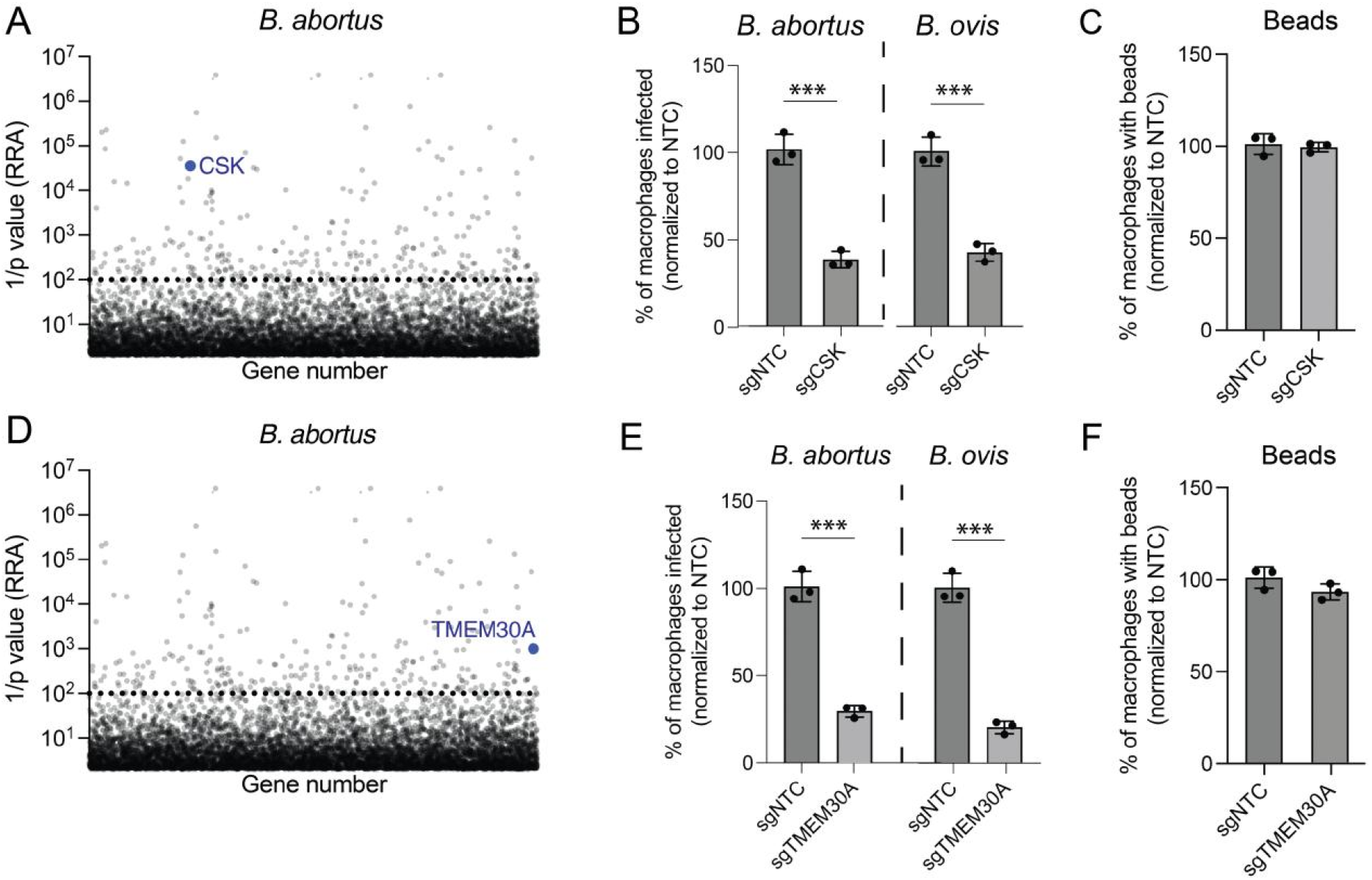
CSK and TMEM30A promote *Brucella* infection of THP-1 macrophages. (A, D) Statistical enrichment plots (1/*P*; dotted line) calculated using the α-RRA algorithm in MAGeCK. CSK (A) and TMEM30A (D) met the criteria for *B. abortus* host factors (≥1 log_2_ fold change, ≥2 independent sgRNAs, *P* < 0.01). (B, E) Validation of CSK (B) and TMEM30A (E) as host factors. THP-1 cells expressing sgRNAs targeting either gene were infected with m*NeonGreen B. ovis* or *B. abortus* at MOI 1,000 for 3 hours. The percentage of infected macrophages was quantified by flow cytometry and normalized to the non-targeting control (NTC). Data represent mean ± SD from three biological replicates. Significance in panels B and E was assessed by an unpaired *t*-test (***, *P* < 0.001). (C, F) Phagocytosis assays using yellow-green fluorescent latex beads (bead:cell ratio = 10) in THP-1 cells expressing NTC or gene-targeting sgRNAs. The percentage of bead-positive cells was quantified by flow cytometry and normalized to NTC. Editing efficiency - CSK: 61%; TMEM30A: 71%.

Several additional cytoskeleton-related genes were identified whose disruption significantly reduced intracellular bacterial burden, including ELMO2, which facilitates actin remodeling during phagocytosis (39). Both CYFIP1 (described above) and WASF2 mutants showed diminished infection in the screen (Table S1). These two genes encode core components of the WAVE complex that drive actin polymerization during dynamic membrane remodeling events (40). Other cytoskeleton/cell motility-related hits included WDR1, an actin disassembly factor that enhances cofilin-mediated filament turnover and cell motility (41). Loss of NHLRC2, which regulates RhoA-Rac1-dependent actin polymerization during phagocytosis (42), also resulted in decreased THP-1 infection. These results are consistent with the known role of actin-driven uptake during *Brucella* entry (3, 43, 44) and expand our view of actin polymerization and disassembly processes that support early stages of macrophage infection by *Brucella*.

### Host lipid metabolism/trafficking as a determinant of Brucella intracellular infection

Our screen also identified several host genes involved in lipid metabolism and trafficking that are required to support *B. abortus* infection in THP-1 cells. Among the top hits in this functional category was TMEM30A, which encodes a subunit of phospholipid flippases responsible for maintaining membrane lipid asymmetry (45) (Fig. 3D). TMEM30A was recently identified as a host factor for *Mycobacterium bovis* BCG (46), SARS-CoV-2 (47, 48), and murine norovirus (49) infection, indicating this gene has an important role in mammalian host cell interactions with both bacterial and viral pathogens. To validate the role of this gene in *Brucella* infection, we generated a TMEM30A mutant THP-1 cell line. Compared to the non-targeting control, the TMEM30A mutant showed significantly reduced intracellular *B. abortus* and *B. ovis* burden (Fig. 3E; Fig. S2C), supporting the screen results. Uptake of fluorescent latex beads was comparable between mutant and control lines (Fig. 3F; Fig. S2D), showing that TMEM30A disruption does not impair general phagocytic capacity.

Several other lipid-related genes whose disruption significantly reduced *B. abortus* infection were identified (Table S1). Among these were PGS1, previously identified as a host factor in an siRNA knockdown screen in HeLa cells infected with *B. abortus* (23), and AGPAT5, both of which participate in ER-localized glycerophospholipid metabolism. Disrupting PTPMT1, a mitochondrial phosphatase required for cardiolipin biosynthesis, DOLK, which produces dolichyl monophosphate for ER-associated glycosylation, and PIP4K2A, a phosphatidylinositol phosphate kinase involved in phosphoinositide signaling and membrane trafficking, also significantly reduced early *B. abortus* infection of THP-1 cells (Table S1). These results provide evidence that host lipid metabolism and homeostasis, in particular ER-localized lipid metabolic processes, play a key role in supporting intracellular *Brucella* infection.

### mTORC1 signaling processes promote B. abortus macrophage infection

Genes categorized as positive regulators of mTOR Complex 1 (TORC1) signaling were significantly enriched among host mutants that showed reduced *B. abortus* infection (Table S1). Although the TORC1 complex itself has not been previously implicated in *Brucella* pathogenesis, the broader mTOR signaling axis is a central regulator of macrophage functions (e.g. polarization, inflammatory signaling, metabolism, and autophagy) that are likely to influence *Brucella* fitness within the host cell environment (50-53). Prompted by this observation, we sought to validate the role of TORC1 signaling-associated genes in promoting early *Brucella* infection. Among the enriched hits, we prioritized LAMTOR2 and AKT1 for functional validation (Fig. 4A; Table S1). Notably, LAMTOR2 was previously identified as a significant hit in an siRNA screen of *B. abortus* infection in HeLa cells (23), providing independent support for its role in *Brucella*-host interactions. THP-1 macrophages with CRISPR-mediated disruptions in either LAMTOR2 or AKT1 were infected with fluorescent *B. abortus* or *B. ovis*, both of which showed significantly reduced intracellular burdens at 3 h compared to non-targeting control cells (Fig. 4B-C; Fig. S3A-B). Again, we tested whether these mutants had a general defect in phagocytosis; latex bead uptake was comparable to control cells in both mutants (Fig. 4D; Fig. S3C).

**Figure 4.**
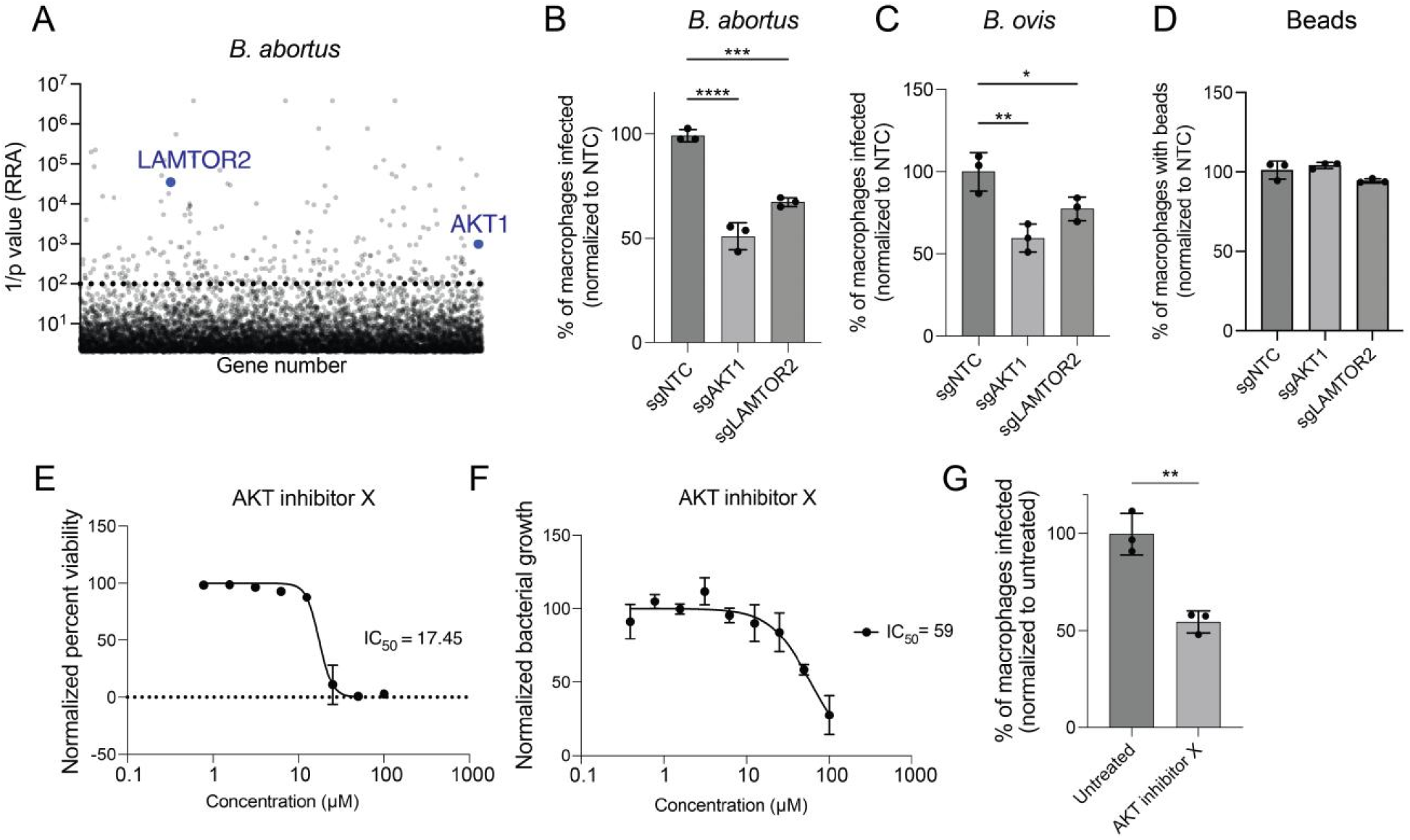
TORC1 signaling genes support efficient *Brucella* infection of macrophages. (A) Plot showing gene enrichment *P* values for the *B. abortus* CRISPR-Cas9 screen, ranked using the α-RRA algorithm in MAGeCK (*P*-value threshold of 10^-2^ (1/p of 10^2^) is drawn as a dotted line). TORC1-related genes *LAMTOR2* and *AKT1* are highlighted. (B-C) Validation of AKT1 and LAMTOR2 as host factors. THP-1 cells expressing sgRNAs targeting either gene were infected with m*NeonGreen B. abortus* (B) or *B. ovis* (C) at MOI 1,000 for 3 hours. The percentage of infected macrophages was quantified by flow cytometry and normalized to the non-targeting control (NTC). Data represent mean ± SD from three biological replicates. Significance in panels B and C was assessed using one-way ANOVA with Dunnett’s multiple comparison test (****, *P* < 0.0001; ***, *P* < 0.001; **, *P* < 0.01). Editing efficiency - AKT1: 63%; LAMTOR2: 87%. (D) Phagocytosis assay in THP-1 macrophages expressing non-targeting control (NTC), *AKT1*, or *LAMTOR2* sgRNAs. Cells were incubated with yellow-green fluorescent latex beads (bead:cell ratio = 10) for 3 hours, and bead-positive cells were quantified by flow cytometry. Data are normalized to the mean uptake in NTC cells. (E) THP-1 viability following treatment with increasing concentrations of AKT inhibitor X for 48 hours, assessed by XTT assay and normalized to untreated controls. (F) Axenic growth of *B. abortus* in broth culture in the presence of AKT inhibitor X. Cultures were incubated for 48 hours, and OD_600_ values were normalized to untreated samples. (G) THP-1 macrophages pretreated with 10 µM AKT inhibitor X showed reduced *B. abortus* infection, as measured by flow cytometry. Data represent mean ± SD from three biological replicates; unpaired *t*-test (*P* < 0.01).

As an orthologous approach to explore the role of AKT in promoting infection, we chemically inhibited AKT activity in THP-1 macrophages using Akt inhibitor X (54). Concentrations above 25 µM impaired host cell viability (Fig. 4E), and doses above 50 µM inhibited *Brucella* growth in axenic culture (Fig. 4F). However, treatment of THP-1 cells with a sub-toxic, sub-inhibitory dose (10 µM) prior to infection significantly reduced intracellular *B. abortus* levels as measured by flow cytometry (Fig. 4G; Fig. S3D), providing further evidence that AKT activity promotes early *Brucella* infection. In addition to LAMTOR2 and AKT1, other significant screen hits with known functional links to TORC1/mTOR signaling included WDFY3 (ALFY), AMBRA1, HDAC3, SLC7A5, DYRK1A, RPTOR, and SIRT6. WDFY3 is reported to localize to RAB5- and EEA1-positive early endosomes in HeLa cells and is highly enriched at dynamic cellular protrusions (55), which may explain its role in supporting early stages of *Brucella* infection. Taken together our results indicate that TORC1 signaling plays an important role in early host-*Brucella* interactions.

### Comparative analysis of host factors in Brucella and Mycobacterium abscessus macrophage infection

To test whether validated host genes that support *Brucella* infection also contribute to infection by an unrelated intracellular pathogen, we challenged the individual mutant THP-1 cell lines with *Mycobacterium abscessus* (Mab), an opportunistic pathogen of increasing concern in people with chronic lung disease (56-58). We infected mutant and non-targeting control THP-1 cells with a previously optimized fluorescent Mab strain (ATCC 19977) that constitutively expresses mEmerald GFP (59) and quantified infection at 4 h post-infection. Disruption of CSK, AKT1, or LAMTOR2 significantly reduced Mab infection relative to controls, whereas RAB14 disruption had no effect (Fig. 5; Fig. S4). Together, these results indicate that CSK, AKT1, and LAMTOR2 support infection by unrelated intracellular pathogens in a THP-1 macrophage model, while RAB14 appears to be *Brucella*-specific.

**Figure 5.**
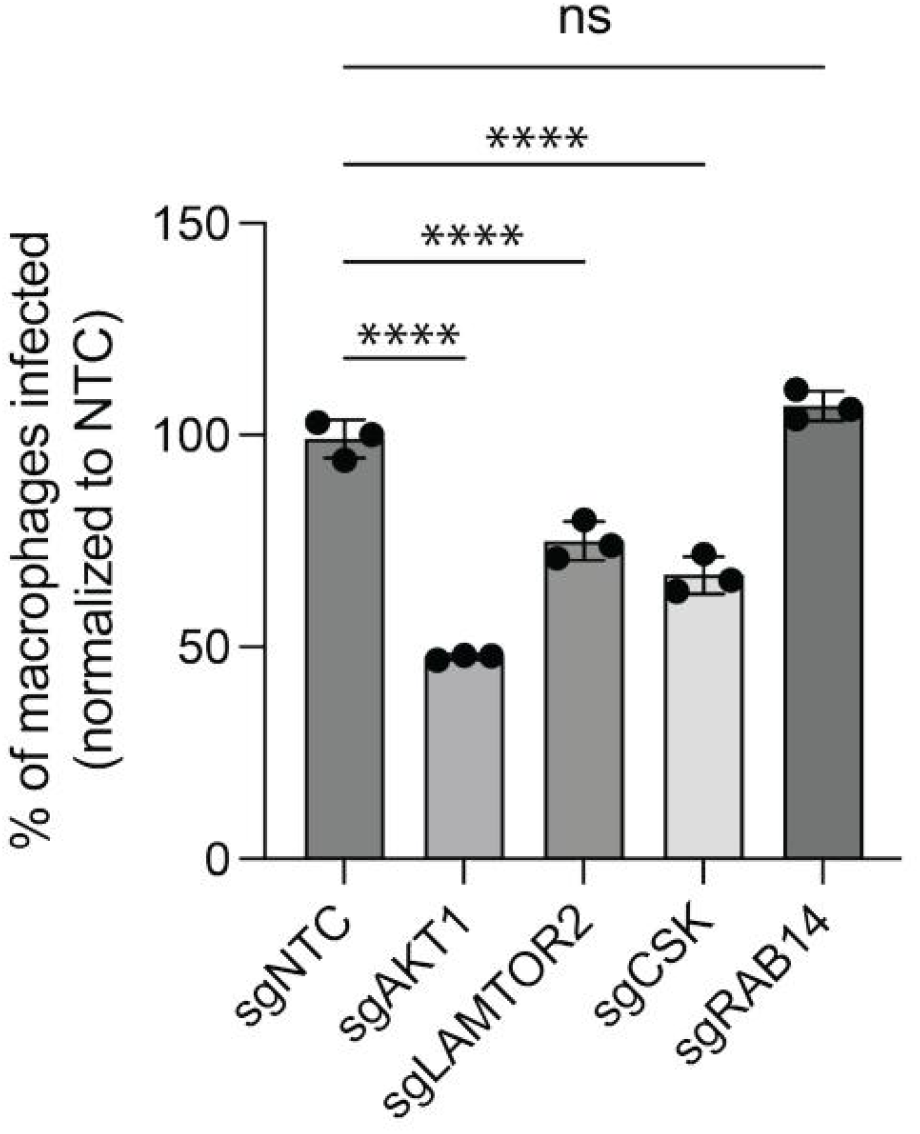
Select THP-1 loss-of-function mutants show reduced susceptibility to *Mycobacterium abscessus* infection. THP-1 macrophages expressing sgRNAs targeting *AKT1, LAMTOR2, CSK*, or *RAB14*, along with a non-targeting control (NTC), were infected with mEmerald-expressing *M. abscessus* at MOI 5 for 4 hours. The percentage of infected cells was quantified by flow cytometry and normalized to NTC. Data represent means ± SD from three biological replicates. Statistical significance was determined using one-way ANOVA followed by Dunnett’s multiple comparison test (****, *P* < 0.0001).

## Discussion

To dissect host-pathogen interactions during the early stages of *Brucella* infection, we developed a flow cytometry-based assay and performed a genome-wide CRISPR-Cas9 loss-of-function screen in human THP-1 cells infected with *B. abortus*. By discriminating infected versus uninfected cells by fluorescence coupled with cell sorting at early time points, we revealed host genes that support the earliest steps of *Brucella* pathogenesis including host entry and the establishment of the endosomal *Brucella*-containing vacuole (eBCV). The identification of known regulators of BCV biogenesis (e.g., CANX, CDC42, RAB5, CYFIP1) (3, 10, 23, 25, 26) in the screen provides evidence that many of the candidates that emerged from our screen are valid. Our follow-up experiments uncovered new genes and pathways that support *Brucella* infection of mammalian hosts, including several pan-infection factors that support infection by *B. abortus, B. ovis*, and an unrelated intracellular pathogen, *M. abscessus*. Thus, our forward genetic screen lays the groundwork for future mechanistic studies of key host pathways discussed below that all contribute to distinct aspects of early host-pathogen interactions.

### Identification of a new Rab GTPase that supports early Brucella infection: RAB14

Consistent with the known role of RAB5 in the formation of nascent *Brucella*-containing vacuoles (BCVs) (25, 26), our screen identified the isoform RAB5C as a host gene that promotes early *B. abortus* infection. The small GTPase RAB14, previously proposed as a possible host factor (60), was also identified as a significant hit. RAB14 coordinates membrane trafficking between early and recycling endosomes and the Golgi, and our results demonstrate that it supports the early stages of infection of THP-1 macrophages by both *B. abortus* and *B. ovis*. In other bacterial and fungal pathogens, including *Mycobacterium tuberculosis* and *Candida albicans*, RAB14 is implicated in delaying endosomal maturation, thereby maintaining a vacuolar environment that favors intracellular persistence (61, 62). Our results raise the possibility that *Brucella* spp. similarly exploit RAB14-regulated pathways to stabilize the eBCV and prevent premature maturation or lysosomal fusion prior to ER remodeling.

RAB14 modulation, however, has pathogen-specific consequences. In *Legionella pneumophila*, RAB14 localizes to the *Legionella*-containing vacuole (LCV) around 1 hour post infection (63), and RAB14 silencing increases intracellular bacterial growth (64). In contrast, *Klebsiella pneumoniae* infection in cells expressing dominant-negative RAB14 shows a significant decrease in intracellular bacterial uptake (65). RAB14 has also been shown to facilitate intracellular replication of *Chlamydia trachomatis* by mediating delivery of sphingolipids to chlamydial inclusions (66). Whether such a sphingolipid delivery mechanism impacts development of the BCV during *Brucella* infection remains an open question. Adding to this repertoire of context-dependent functions, we found that RAB14 was dispensable for infection by *Mycobacterium abscessus*, indicating that RAB14 is not universally required for intracellular bacterial growth or restriction. Together, our data and prior work suggest that RAB14 differentially interacts with intracellular bacteria to either enhance or restrict uptake and growth. This specificity almost certainly reflects important differences in intracellular trafficking dynamics between *Brucella* spp., *M. abscessus*, and other bacterial species.

The absence of late endosomal and lysosomal markers such as RAB7 and LAMP1 among our top candidates supports the idea that our 3 hour post-infection screening window is enriched for host factors involved in the earliest stages of infection (specifically, eBCV formation and early trafficking events). Future studies that track RAB14 dynamics on BCVs and dissect its effector networks will clarify whether RAB14 primarily sustains a pro-entry niche or plays other roles in modulating BCV interaction with other compartments.

### CSK as a conserved determinant of early infection

CSK is a negative regulator of Src-family kinases (SFKs), that maintains SFKs in an inactive conformation via C-terminal phosphorylation (67). SFKs, in turn, influence multiple processes relevant to pathogen entry, including actin remodeling, endocytic routing, and inflammatory signaling. Our discovery that CSK disruption reduces early infection by *B. abortus, B. ovis*, and *M. abscessus* suggests that precise SFK tuning by CSK favors successful establishment of intracellular niches for both *Brucella* and *M. abscessus*. This conclusion is consistent with prior work linking CSK to microbial entry phenotypes. For example, CSK perturbation has been associated with reduced internalization of *Pseudomonas aeruginosa* (38) and with effects on viral replication (37).

CSK-deficient THP-1 cells retained normal uptake of fluorescent beads, arguing against a global defect in phagocytosis and instead pointing to signaling abnormalities that specifically affect BCV establishment. Given the known roles of CSK and SFKs in host cell biology, defective coordination of early endosomal trafficking remains a likely mechanism underlying the attenuated infection observed in CSK mutant lines (68, 69). One possibility is that loss of CSK derepresses SFK activity in a manner that promotes bactericidal outcomes, for example by enhancing endosome-lysosome fusion, but this remains to be tested experimentally.

### On TORC1 signaling and Brucella infection

GSEA implicated mTORC1-related processes as important for the early stages of *Brucella* infection (Table S1); genes with connections to TORC1 signaling including WDFY3, AMBRA1, HDAC3, SLC7A5, DYRK1A, RPTOR, and SIRT6 emerged as significant host factor hits in our screen. AKT–mTORC1 signaling is a central regulator of macrophage metabolism, polarization, and inflammatory programs (50, 53, 70, 71), all of which can condition the intracellular niche. Consistent with this, pharmacologic inhibition of AKT1 at a non-toxic, sub-host inhibitory dose reduced early *B. abortus* infection, mirroring the reduction in intracellular burden observed in AKT1-deficient THP-1 cells and aligning with prior work showing that select intracellular bacteria and viruses exploit AKT signaling to support survival or replication (72, 73). AKT1-mutant THP-1 cells were also less susceptible to *M. abscessus* infection, suggesting that AKT1-dependent signaling promotes the establishment of a permissive environment for disparate intracellular bacterial pathogens. It remains an open question whether AKT1 helps to promote a Warburg-like metabolic state that was previously reported to support *Brucella* infection (74), or whether signaling differences between AKT isoforms (AKT1 versus AKT2) differentially shape macrophage inflammatory responses during infection (75).

In addition to AKT1, we validated LAMTOR2 as a host factor for *B. abortus, B. ovis*, and *M. abscessus* infection of THP-1 macrophages. This protein is a core component of the Ragulator complex (76), where it coordinates amino acid sensing with ERK/MAPK signaling (77, 78), influencing both mTORC1 activation and lysosomal function (79). Notably, one of the top hits in our screen, DYRK1A, was recently shown to interact with the TSC1-TSC2 complex where it phosphorylates TSC2 to promote TORC1 activation (80). Thus, loss of DYRK1A function may reduce *Brucella* infection by dampening TSC–mTORC1 signaling in macrophages. Consistent with a nutrient-sensing contribution to this signaling axis, we also identified SLC7A5 (LAT1) as a host factor (Table S1). This large neutral amino acid transporter mediates leucine uptake and is required for full mTORC1 activation and metabolic reprogramming in immune cells (81-83). Altogether, these results point to a broad network of regulators that converge on mTORC1 to influence a *Brucella*-permissive intracellular state.

Contrasting with our results, previous infection studies in LAMTOR2^−/−^ macrophages reported increased intracellular bacterial loads 6–24 h after infection with *Salmonella enterica* serovar Typhimurium or *Klebsiella pneumoniae* (84, 85). In those systems, LAMTOR2 thus appears to limit intracellular replication. Our result that LAMTOR2 disruption reduces early infection by two *Brucella* species and by *M. abscessus*, indicates that LAMTOR2 can act as either a pro-infection host factor or a restriction factor depending on the organism, cell type, and stage of infection.

It is possible that *Brucella* depends on Ragulator-dependent lysosomal cues during eBCV maturation (e.g. controlled changes in luminal pH, ion composition, or positioning of BCVs within the endolysosomal network) that are required to trigger *virB* (Type IV secretion system) expression and subsequent ER remodeling without committing to terminal lysosomal fusion. In this model, loss of LAMTOR2 and other positive regulators of TORC1 signaling could misregulate these cues, diverting BCVs into more bactericidal pathways. Dissecting if and how Ragulator–mTORC1 and AKT1 integrate metabolic, lysosomal, and signaling inputs with other validated host factors at early timepoints will be essential to understand why mTORC1 pathway components are required for efficient infection by *Brucella* and *M. abscessus*. Altogether, our functional genetic approach identified and validated distinct host cell processes that are required for *Brucella* infection and may be potential host targets to limit diverse intracellular pathogens.

## Materials and Methods

### Bacterial strains and growth conditions

*Brucella ovis* 25840 and *Brucella abortus* 2308 were grown on tryptic soy agar (TSA; Difco Laboratories) plates supplemented with 5% (v/v) sheep blood (Quad Five) or in brucella broth (BB; Difco Laboratories) dissolved in milliQ water for liquid cultures. *B. ovis* and *B. abortus* cells were incubated at 37ºC with 5% (v/v) CO_2_ supplementation. Smooth Mab (ATCC 19977) cultures were grown aerobically in Middlebrook 7H9 medium supplemented with 10% Middlebrook OADC (oleic acid, dextrose, catalase, and bovine albumin) at 37ºC. mEmerald GFP-expressing strains were propagated in medium containing 5 µg/mL zeocin (Invivogen). All *Escherichia coli* strains were grown in liquid LB or on LB solidified with 1.5% (w/v) agar. Stbl3 or Top10 strains were incubated at 30ºC or 37ºC respectively. Strain information is available in Table S2.

### Construction of Brucella mNeonGreen strains

A *B. abortus* strain that constitutively expresses mNeonGreen fluorescent protein was generated from the wild-type *B. abortus* 2308 parent strain by integration of a pUC18-mTn7-kanamycin plasmid harboring mNeonGreen at the *glmS* locus (the mini-Tn7 plasmid was gift from H.P Schweizer). The pUC18-mTn7-mNeonGreen-kanamycin plasmid was co-conjugated into *B. abortus* with the helper plasmid pTNS3 expressing the Tn7 integrase gene as previously described for construction of lux-expressing *Brucella* strains (86). *B. abortus* colonies carrying the integrated mTn7-mNeonGreen construct were selected on TSA blood plates containing 50 µg ml^-1^ kanamycin.

### THP-1 cell culture

THP-1 macrophage-like cells were grown to a maximum titer of 1 x 10^6^ cells ml^-1^ in complete RPMI 1640 medium supplemented with 2 mM glutamine (GIBCO), and 10% heat-inactivated fetal bovine serum (v/v) (HI FBS) (HyClone).

### THP-1 infections and treatments

#### THP-1 Brucella infection

THP-1 cells were seeded at a titer of 1 x 10^6^ cells per well in 6-well plates, and phorbol myristate acetate (PMA) was added at a final concentration of 50 ng µl^-1^ to induce differentiation into macrophage-like cells for 48-72 h prior to infection. *B. ovis* and *B. abortus* strains expressing mNeonGreen were harvested from 48-hour-old plates, resuspended in RPMI containing 10% HI FBS, and added to tissue culture plates on the day of infection at a multiplicity of infection (MOI) of 1000. Plates were centrifuged for 5 min at 150 x g and incubated for 3 h at 37ºC in 5% (v/v) CO_2_. The medium was removed, and fresh medium containing 50 µg ml^-1^ gentamicin was supplied, followed by a 30-minute incubation. Following gentamicin incubation, the medium was removed, and cells were washed with PBS. Cells were treated with TrypLE Express (Gibco) at 37ºC for 15 minutes and gently lifted and harvested. Harvested cells were centrifuged at 500 × g and washed once with PBS. For viability analysis, the supernatant was aspirated and cells were incubated for 15 minutes at room temperature in the dark with Zombie NIR Fixable Viability Dye (BioLegend) diluted in PBS. Stained cells were washed with Cell Staining Buffer (BioLegend), fixed in 3.2% paraformaldehyde for 20 minutes at room temperature, and washed three times with PBS. After completing a confirm-kill protocol, macrophage infection was quantified using the Attune CytPix benchtop analyzer (ThermoFisher).

#### THP-1 chemical pretreatment

THP-1 cells were seeded at 1 × 10^6^ cells per well in 6-well plates and treated with 50 ng/µL PMA to induce differentiation into macrophage-like cells for 48–72 hours prior to infection. Twenty-four hours before infection, AKT inhibitor X (Cayman Chemical) was added to the culture media at a final concentration of 10 µM and incubated at 37°C with 5% CO_2_. On the day of infection, mNeonGreen-expressing *Brucella abortus* cells were resuspended from 48-hour-old plates in RPMI supplemented with 10% (v/v) heat-inactivated FBS. Media from plates were removed and replaced with fresh media containing *B. abortus* at a multiplicity of infection (MOI) of 1,000. Plates were centrifuged at 150 × g for 5 minutes, then incubated for 3 hours at 37 °C with 5% CO_2_. Following infection, media were removed, cells were washed with PBS, and macrophages were detached using TrypLE Express (Gibco) at 37°C for 15 minutes. Lifted cells were gently harvested, centrifuged at 500 × g, and washed once with PBS. Cells were then fixed in 3.2% paraformaldehyde for 20 minutes at room temperature, followed by three PBS washes. After completing a confirm-kill protocol, macrophage infection was quantified using the Attune CytPix benchtop analyzer (ThermoFisher). Cell viability was assessed as described above.

#### THP-1 Mycobacterium abscessus (Mab) infection

THP-1 cells were seeded at 1 × 10^6^ cells per well in 6-well plates and treated with 50 ng/µL PMA for 48–72 hours to induce differentiation into macrophage-like cells, as described above. Log-phase *M. abscessus* cultures were resuspended in RPMI supplemented with 10% (v/v) heat-inactivated FBS, and large clumps were removed by centrifugation at 800 × g for 5 minutes. The resulting single-cell suspension was used to infect THP-1 macrophages at a multiplicity of infection (MOI) of 5. Infected plates were centrifuged at 150 × g for 5 minutes to promote contact and incubated for 4 hours at 37°C with 5% CO_2_. Following infection, media were removed, and cells were washed with PBS. Macrophages were detached using TrypLE Express (Gibco) at 37 °C for 15 minutes, then gently harvested, centrifuged at 500 × g, and washed once with PBS. For viability assessment, cells were incubated with Zombie NIR fixable viability dye (BioLegend) diluted in PBS for 15 minutes at room temperature in the dark, followed by a wash with cell staining buffer (BioLegend). Cells were then fixed in 3.2% paraformaldehyde for 20 minutes and washed three times with PBS. After completing a confirm-kill protocol, infection and viability were quantified using the Attune CytPix benchtop analyzer (ThermoFisher).

#### Fluorescent bead uptake assay

1 µm carboxylate-modified yellow-green fluorescent polystyrene latex beads (Sigma-Aldrich) were diluted in PBS to 1 x 10^7^ beads per ml. Beads were incubated with THP-1 cells for 3 h. Following bead exposure, macrophages were washed with PBS and fixed in 3.2% paraformaldehyde solution for 20 minutes. The percentage of macrophages with yellow-green fluorescence latex beads was quantified by flow cytometry using the Attune CytPix benchtop analyzer.

### CRISPR screen and analysis

We used the Human Brunello CRISPR knockout pooled guide library (Addgene #73178; gift of David Root and John Doench) (24) to generate a THP-1 gene knockout pool. Briefly, lentiviral particles were produced and used to transduce THP-1 cells stably expressing Cas9 at low MOI to favor single-sgRNA integration. Over 99.2% of sgRNAs were detected in the resulting mutant pool used for infection. The pool had a low Gini index (0.091) indicating good evenness across the mutant library (average mapped read depth of ∼540 reads/sgRNA).

This THP-1 mutant library was infected with a *B. abortus* strain that constitutively expressed mNeonGreen from a Tn7 construct integrated at the *glmS* locus; infections were at an MOI of 1,000 for 3 h. After 3 h, macrophages were washed with PBS then fixed with 3.2% paraformaldehyde. The THP-1 cells were then sorted in BSL3 containment using a Bio-Rad S3e cell sorter to isolate mNeonGreen+ (i.e. infected) and mNeonGreen-(i.e. uninfected) macrophages. A total of 1.5-2.5 x 10^7^ cells were sorted into each bin from two replicate samples. Genomic DNA was isolated from each sorted population using Qiagen DNeasy kits after reversing DNA cross-links following a 55°C incubation overnight. Amplification of sgRNAs by PCR was performed as previously described (87) using Illumina compatible primers from IDT and amplicons were sequenced on an Illumina NovaSeq (*B. abortus* reads). Amplicon sequencing depth was assessed for the *B. abortus* experiment (∼15 million mapped reads per sample). Sequenced reads were trimmed to remove any adapter sequence and to adjust for the p5 primer stagger. We used model-based analysis of genome-wide CRISPR-Cas9 knockout (MAGeCK) version 0.9.5.0 (88) to count reads with matches to the Brunello sgRNA library index without allowing for any mismatch. Subsequent sgRNA counts were median normalized in MAGeCK to account for variable sequencing depth. To test for sgRNA and gene enrichment, we used the test command in MAGeCK to compare the distribution of sgRNAs in the mNeonGreen+ and mNeonGreen-bins. To identify host factors that influence *Brucella* infection, we used the following criteria: 1) detection of at least two sgRNAs in the dataset, 2) an experimental log selection ratio greater than 1, and 3) experimental *P* value less than 0.01.

### Gene set enrichment analysis

Gene set enrichment analysis (GSEA) was performed in R v4.4.3 using the packages fgsea and msigdbr. All pathway-level GSEA analyses of host factors were based on the *B. abortus* screen dataset. Gene-level statistics were imported from a tab-delimited file containing gene symbol, *p* value, and log_2_ fold change. To generate a directional pre-ranked list, we computed a gene score defined as (−log_10_ *p* value) × (−log_2_ fold change), so that genes whose disruption most strongly depleted infection (large negative (Neg LogFC) and small *p* value) received large positive scores. Open gene sets for *Homo sapiens* were obtained from MSigDB via msigdbr, using the Gene Ontology Biological Process collection (C5, GO:BP) and Reactome canonical pathways (C2, REACTOME). For each collection, we required gene sets to contain between 15 and 500 genes. One-sided multilevel GSEA was then performed with `fgseaMultilevel`, testing the positive tail of the ranked list (“pos”; enrichment among host factors) and the negative tail (“neg”; enrichment among putative restriction factors) separately for each collection. This procedure returns, for every gene set, a normalized enrichment score (NES), nominal *p* value, Benjamini-Hochberg-adjusted *p* value (FDR), gene set size, and leading-edge subset. Results from all four analyses (GO:BP-pos, GO:BP-neg, Reactome-pos, Reactome-neg) were combined into summary tables and filtered by FDR. Using an arbitrary significance threshold of adjusted *p* value (FDR) < 0.1, no gene sets were significantly enriched in the negative (“restriction factor”) tail; thus only host factor-enriched categories with at least one leading-edge gene (CORE_ENRICHMENT = TRUE) are reported in Table S1.

### Generation of targeted CRISPR knockout lines

Individual sgRNAs were cloned into the sgOPTI vector (Addgene plasmid #85681, from the Eric Lander lab) as previously described (87). Briefly, annealed sgRNA oligonucleotides were phosphorylated and ligated into BsmBI-digested, dephosphorylated sgOPTI-Puromycin-Ampicillin (PA) plasmids. Transformants were selected on ampicillin-containing plates, and at least one sgRNA construct per target gene was packaged into lentivirus for transduction into Cas9-expressing THP-1 cells. Following puromycin selection, genomic DNA was extracted from transductants. Target regions were PCR-amplified and submitted to the Michigan State University Genomics Core for Sanger sequencing. Editing efficiency and indel size were quantified from the Sanger sequence data using Tracking of Indels by Decomposition (TIDE) analysis (89). For each gene, a single knockout line with >60% editing efficiency was selected for downstream experiments. sgRNA sequences and TIDE primers are listed in Table S3.

### Colorimetric THP1 XTT cell proliferation assay

THP-1 cells were seeded at a titer of 1 x 10^5^ cells per well in RPMI supplemented with 10% (v/v) FBS and containing 50 ng µl^-1^ PMA to induce differentiation into macrophage-like cells in 96-well plates (Corning) for 48–72 h. Varying concentrations of compounds were prepared by hand and transferred to plates containing THP1 cells with final concentrations of each compound (suspended in RPMI and 10% [v/v] FBS) ranging from 1 µM to 100 µM. After incubation, 50 µl XTT (sodium 3’-[1-(phenyl aminocarbonyl)-3,4-tetrazolium]-bis (4-methoxy-6-nitro) benzene sulfonic acid hydrate) (Roche) was added to each well and incubated for 4 h at 37ºC and 5% (v/v) CO^2^. Following incubation, absorbance of the reaction product was measured at 570 nm and 650 nm.

## Acknowledgements

We thank Lydia Varesio for the construction of the pUC18 glmS-mTn7-mNeonGreen plasmid. We also thank the Olive lab for assistance with Mab cultivation and infection and the MSU Flow Cytometry Core for instrumentation support. Research reported in this publication was supported in part by the NIH award number R01AI177619 to S.C. and R35GM146795 to A.O.

## Supplementary Figures

**Figure S1.**
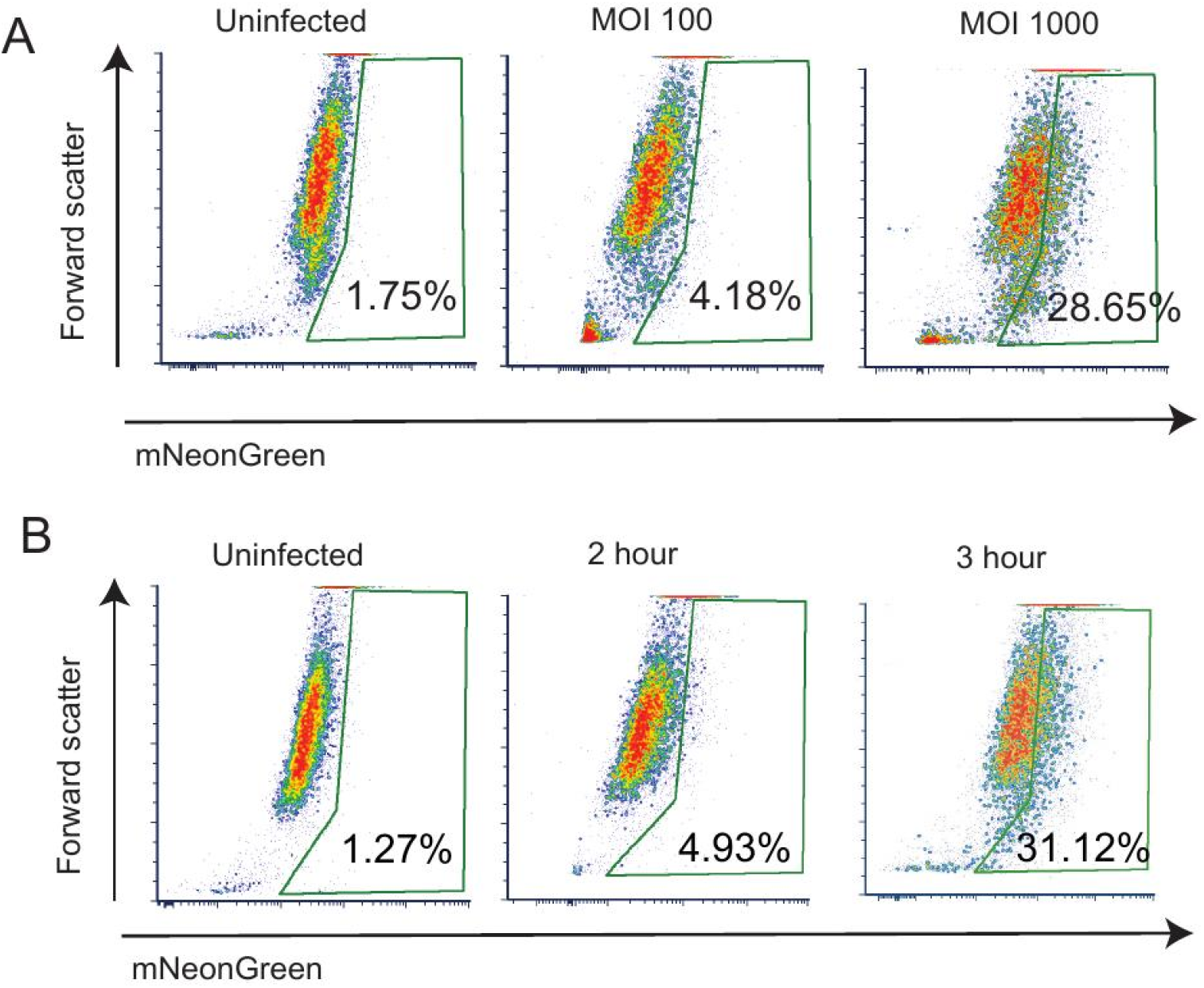
*B. ovis* efficiently infects THP-1 macrophages at 3 hours post-infection at a multiplicity of infection (MOI) of 1,000. (A) Representative flow cytometry plots showing mNeonGreen-positive THP-1 macrophages at 3 hours post-infection at increasing MOI. Plots are gated on live, single cells (gate shown as green box on flow plots) and the percentage of mNeonGreen-positive (infected) cells is indicated in each plot. (B) Representative flow cytometry plots showing mNeonGreen-positive macrophages at 2 and 3 hours post-infection with *B. ovis* at MOI 1,000. All plots are gated on live, single cells.

**Figure S2.**
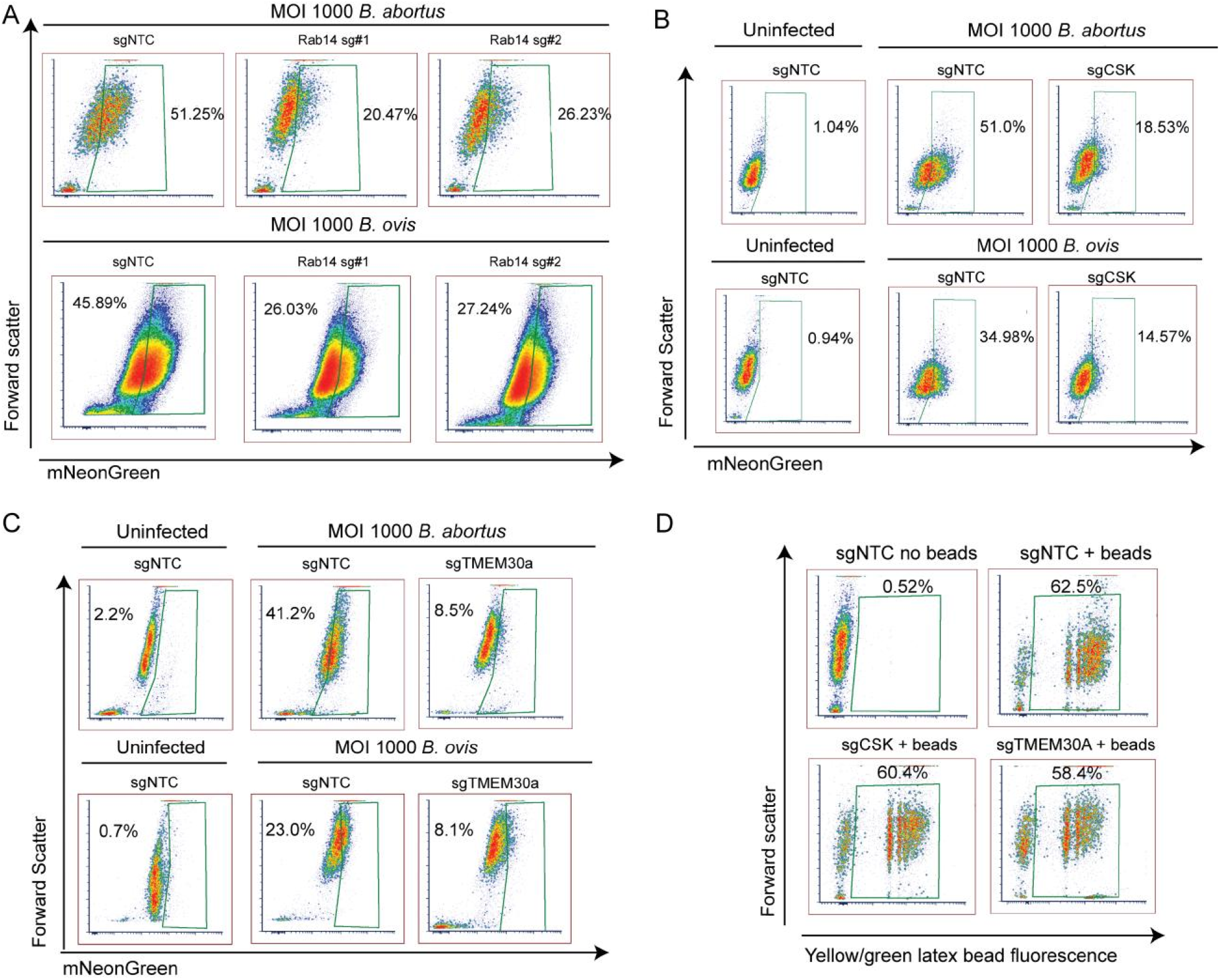
RAB14-, CSK-, and TMEM30A-deficient THP-1 macrophages are less susceptible to infection by *B. abortus* and *B. ovis*. THP-1 macrophages expressing non-targeting control (sgNTC) or the indicated sgRNAs were infected for 3 h with mNeonGreen-expressing *B. abortus* or *B. ovis* at MOI 1,000. Live, single cells were gated (gate shown as green box on flow plots), and the percentage of mNeonGreen-positive (infected) cells is indicated in each plot. (A) Representative plots for sgNTC cells and two independent RAB14-targeting sgRNA lines infected with *B. abortus* (top row) or *B. ovis* (bottom row). (B) sgNTC and CSK-deficient (sgCSK) cells, shown uninfected (left panels) or infected with *B. abortus* (top row) or *B. ovis* (bottom row). (C) sgNTC and TMEM30A-deficient (sgTMEM30A) cells, with uninfected sgNTC controls (left panels) and infections with *B. abortus* (top row) or *B. ovis* (bottom row). (D) Latex bead uptake in sgNTC, CSK-deficient, and TMEM30A-deficient macrophages. Cells were incubated with yellow-green fluorescent beads (bead:cell ratio 10:1) for 3 h. Representative plots show similar proportions of bead-positive cells across genotypes, indicating that phagocytic capacity is preserved in the mutant THP-1 lines.

**Figure S3.**
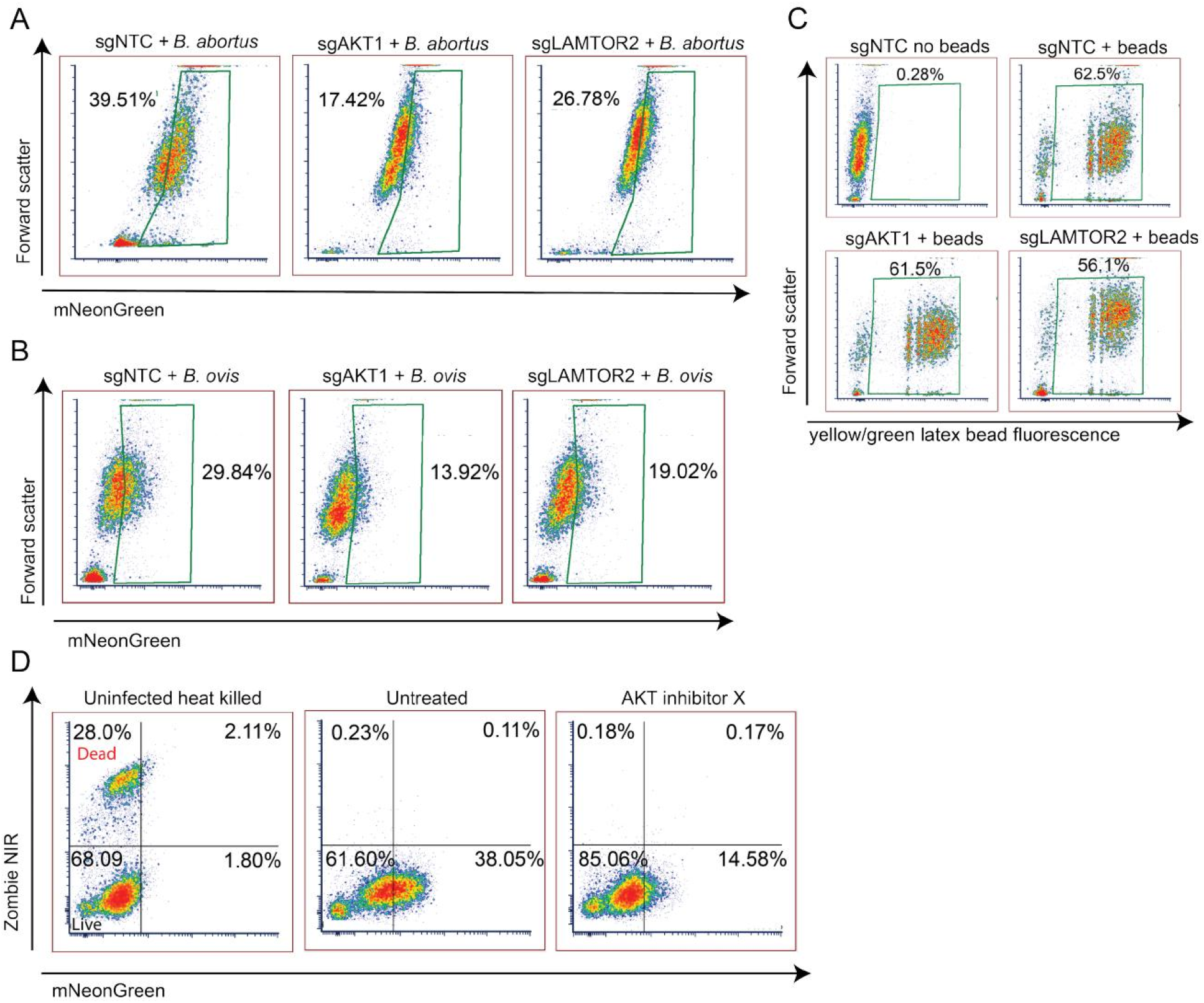
AKT1- and LAMTOR2-deficient THP-1 macrophages are less susceptible to infection by *B. abortus* and *B. ovis*, and pharmacological AKT inhibition reduces *B. abortus* infection without inducing cytotoxicity. THP-1 macrophages expressing non-targeting control (sgNTC) or sgRNAs targeting AKT1 or LAMTOR2 were infected with mNeonGreen-expressing *B. abortus* or *B. ovis* (MOI = 1,000) and analyzed 3 h post-infection by flow cytometry. (A, B) Representative flow cytometry plots showing the percentage of infected (mNeonGreen-positive) cells within the live, single-cell gate for each genotype after infection with *B. abortus* (A) or *B. ovis* (B). (C) Latex bead uptake in sgNTC, AKT1-deficient, and LAMTOR2-deficient macrophages. Cells were incubated with yellow-green fluorescent beads (bead:cell ratio 10:1) for 3 h. Representative plots show similar proportions of bead-positive cells across genotypes, indicating that phagocytic capacity is preserved in the mutant THP-1 lines. (D) Representative viability and infection profiles of THP-1 macrophages 3 h after infection with mNeonGreen *B. abortus* under the indicated conditions. Cells were gated (horizontal and vertical lines on the flow plot) on live, single events and stained with Zombie NIR viability dye. The left panel shows uninfected, heat-treated macrophages used to define live/dead gates. The middle and right panels show macrophages that were either untreated or pre-treated with 10 micromolar AKT inhibitor X prior to infection. Quadrants indicate the percentages of live/dead (Zombie NIR) and infected (mNeonGreen-positive) cells.

**Figure S4.**
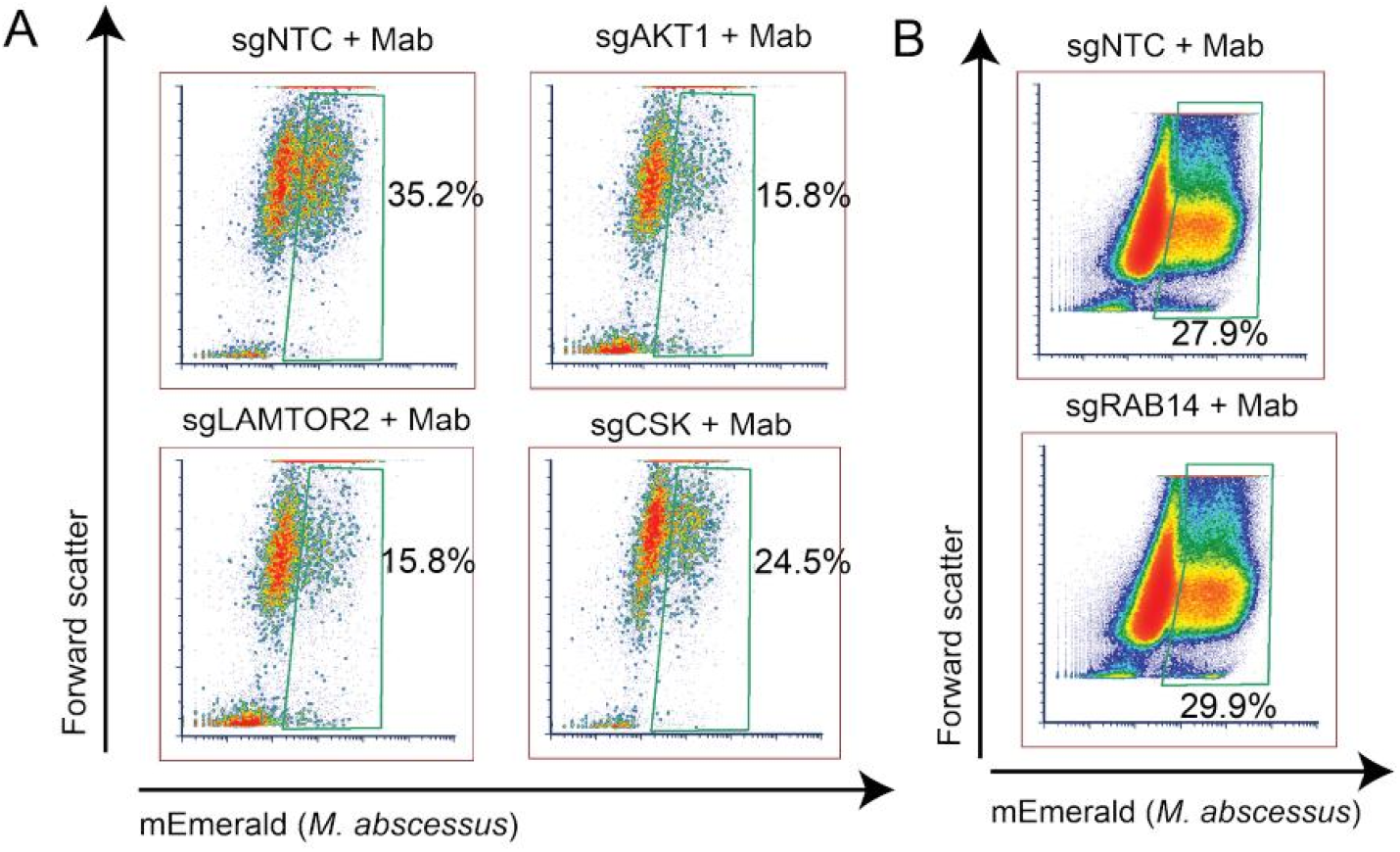
*Brucella* host factors also support *M. abscessus* infection, with the exception of RAB14. (A) Flow cytometry plots showing THP-1 macrophages expressing non-targeting control (sgNTC) or sgRNAs targeting AKT1, LAMTOR2, or CSK, infected with *mNeonEmerald*-expressing *Mycobacterium abscessus* at MOI = 5 for 4 hours. The percentage of mEmerald-positive (infected) cells is indicated for each genotype within the live, single-cell gate (gate shown as green box in the flow plots). (B) Representative flow cytometry plots for THP-1 cells expressing sgNTC or sgRNAs targeting RAB14, infected with *M. abscessus* under the same conditions. Unlike other validated *Brucella* host factors, disruption of RAB14 did not reduce *M. abscessus* infection, indicating a *Brucella*-specific requirement for this gene.

## Notes

### Competing Interest Statement

The authors have declared no competing interest.

